# SARS-CoV-2 infects neurons, astrocytes, choroid plexus epithelial cells and pericytes of the human central nervous system

**DOI:** 10.1101/2023.11.21.568132

**Authors:** Ruth Haverty, Janet McCormack, Christopher Evans, Kevin Purves, Sophie O’Reilly, Virginie Gautier, Keith Rochfort, Aurelie Fabre, Nicola F. Fletcher

## Abstract

SARS-CoV-2, the coronavirus responsible for the COVID-19 pandemic, is associated with a range of neurological manifestations including haemorrhage, thrombosis and ischaemic necrosis and encephalitits. However, the mechanism by which this occurs is unclear. Neurological disease associated with SARS-CoV-2 infection has been proposed to occur following direct infection of the central nervous system and/or indirect sequelae as a result of peripheral inflammation. We profiled ACE2 and TMPRSS2 in brain tissue from five healthy human donors, and observed expression of these proteins in astrocytes, neurons and choroid plexus epithelium within frontal cortex and medulla. Primary human astrocytes, neurons and choroid plexus epithelial cells supported productive SARS-CoV-2 infection in an ACE2- dependent manner. Infected cells supported the full viral lifecycle, releasing infectious virus particles. In contrast, primary brain microvascular endothelial cells, pericytes and microglia were refractory to SARS-CoV-2 infection. These data support a model whereby SARS-CoV-2 is neurotropic, and this may in part explain the neurological sequelae of infection.

**Importance:** A subset of patients with COVID-19 develop neurological symptoms, but the underlying cause is poorly understood. We observed that cells within normal human brain express the SARS-CoV-2 entry factors ACE-2 and TMPRRS2, with expression mainly observed within astrocytes, neurons and choroid plexus epithelium. Primary human astrocytes, neurons and choroid plexus epithelial cells cultured *in vitro* supported the full SARS-CoV-2 life cycle with a range of SARS-CoV-2 variants. This study demonstrates that cells of the human central nervous system express SARS-CoV-2 entry factors *in vivo* and support viral infection *in vitro*, thus supporting a model where neurological symptoms seen in some COVID-19 patients may be as a result of direct viral infection of the central nervous system. Furthermore, these data highlight the importance of investigating the ability of therapeutics to clear virus from this potential reservoir of infection.

## 1. Introduction

SARS-CoV-2, a member of the Coronaviridae, is a positive-sense, single-stranded RNA virus with a genome size of ∼30 kb (1). Responsible for the COVID-19 pandemic, it has caused approximately 7 million confirmed deaths (2). While infection is primarily associated with respiratory disease (1), chronic complications are increasingly recognized in recovered patients (3, 4). These include neurological sequelae including fatigue and cognitive dysfunction, insomnia, gastrointestinal symptoms, arthralgia and other chronic symptoms, collectively referred to as post-acute sequelae of SARS-CoV-2 infection (PASC)(4). In addition, neurological symptoms including loss of smell and taste, are observed in acutely infected individuals (4). The pathogenesis of SARS-CoV-2 neurological disease is incompletely understood, and may be due to viral tropism for cells of the central and peripheral nervous systems and/or as a result of peripheral inflammation or other mechanisms (4).

A number of studies have investigated whether SARS-CoV-2 can directly infect the central nervous system (CNS). Recent studies have reported viral antigen in choroid plexus organoids with disruption of the blood-cerebrospinal fluid (CSF) barrier (3, 5), and productive viral infection of neural progenitor cells and brain organoids (6). In contrast, Pedrosa et al. did not detect SARS-CoV-2 infection of human neurospheres (7), although an inflammatory response was observed following exposure to virus. *In vivo*, SARS-CoV-2-like particles have been detected in brain microvascular endothelium (8). A further study investigating SARS-CoV-2 infection of neural organoids together with mouse and human brain detected viral antigen in neuronal cells and cortical neurons in post-mortem human brain tissue (9). Therefore, while there is increasing evidence that SARS-CoV-2 is neurotropic, there remain questions on the main cell types within the CNS that support viral infection, the role of ACE-2 and other receptors in viral infection and the ability of infected cells to support the full viral lifecycle remains unclear.

We investigated SARS-CoV-2 receptor expression and SARS-CoV-2 tropism in post-mortem human brain tissue and a panel of primary cells isolated from human brain tissue, including primary brain microvascular endothelial cells, astrocytes, neurons, pericytes, choroid plexus epithelial cells and microglia. The ability of permissive cells to support the full viral lifecycle and release infectious viral particles was investigated, together with the ability of an ACE-2 neutralising antibody to inhibit SARS-CoV-2 infection. Finally, the ability of permissive cells to support infection with alpha, delta and omicron BA.1 variants was investigated. These data have implications for development of therapeutics against SARS-CoV-2, which may require delivery to the CNS across the blood-brain barrier, and provide novel information on the neuropathogenesis of COVID-19-associated neurological disease.

## 2. Methods

### 2.1 Immunohistochemistry to determine ACE2 and TMPRSS2 expression in normal human brain tissue

Normal human brain tissue was obtained from the MRC Edinburgh Brain & Tissue Bank (MRC Database Number BBN). Tissue was donated with consent according to the Human Tissue (Scotland) Act with ethical approval number 16/ES/0084 and local ethical exemption from University College Dublin (UCD Human Research Ethics Committee (HREC) Approval No. HREC-LS-20-358469). Sequential 4μm sections of formalin-fixed, paraffin-embedded normal brain tissue from 5 donors were dewaxed, rehydrated and antigen retrieval was performed in a PTLink with EDTA buffer (TMPRSS2 ab109131, Abcam) and citrate buffer (ACE2 ab15348, Abcam) and stained on a AS Link 48 autostainer. After incubation with primary antibodies (0.8 μg/mL (TMPRSS2), 1 μg/mL (ACE2)), sections were incubated with an HRP-polymer, revelated using diaminobenzidine (DAB) chromogen and counterstained with haematoxylin (EnVision Flex HRP High pH Kit, K8002, Agilent Technologies). The stained slides were scanned at 20X magnification using the Aperio AT2 digital slide scanner (Leica Microsystems). Slides were evaluated for positive TMPRSS2 and ACE2 staining by a board certified pathologist (AF).

### 2.2 Cell lines, primary cells and antibodies

VeroE6 cells (ATCC CRL-1586) were propagated in Dulbecco’s modified Eagle’s medium (DMEM) + GlutaMAX™-I (Gibco) supplemented with 10% Foetal bovine serum (FBS) and 1% non-essential amino acids (Gibco). Primary human neurons (HN; 1520-10), primary human astrocytes (HA; 1800-10), primary human choroid plexus epithelial cells (HCPEpiC; 1310) and primary human brain vascular pericytes (HBVP; 1200) were obtained from ScienCell (Carlsbad, CA, USA). Primary cells were cultured as indicated by ScienCell, and maintained in Neuronal Medium supplemented with 1% Neuronal Growth Supplement (HN), Astrocyte Medium supplemented with 1% Astrocyte Growth Supplement (HA), Epithelial Cell Medium supplemented with 1% Epithelial Cell Growth Supplement (HCPEpiC) or Pericyte Medium supplemented with 1% Pericyte Growth Supplement (HBVP), together with 2% FBS (ScienCell) and 1% Penicillin/Streptomycin (ScienCell). Primary Human Brain Microvascular Endothelial Cells (BMEC; ACBRI 376) were obtained from Cell Systems (Kirkland, WA, USA) and maintained in Serum-containing Basal Medium (Cell Systems) supplemented with 1% CultureBoost™ (Cell Systems), and 0.05% Bac-Off® (Cell Systems). Immortalised Human Microglia (IHM; P10354-IM) were obtained from Innoprot (Bizkaia, Spain) and cultured in Microglial Basal Medium (Innoprot) supplemented with 1% Microglial growth supplement (Innoprot), 5% FBS and 1% penicillin/streptomycin. All cells were maintained at 37°C, 5% CO_2_.

Primary antibodies used for immunofluorescent staining and neutralisation assays were as follows: SARS-CoV-2 Spike mAb anti-rabbit IgG (SinoBiological, Beijing, China); dsRNA K1 mAb anti-mouse IgG2a (Scicons, Susteren, The Netherlands), and recombinant anti-ACE2 neutralising mAb anti-mouse IgG1 (SinoBiological, Beijing, China). Fluorescent secondary antibodies Alexa Fluor 488 and 594 goat anti-rabbit and anti-mouse, respectively, were obtained from Invitrogen.

### 2.3 Virus culture and isolates

SARS-CoV-2 (2019-nCoV/Italy-INMI1) obtained from the European Virus Archive Global (EVAg; GenBank accession MT077125.1), was propagated in VeroE6 cells. All virus used in this study was at passage 3. Cells were inoculated with an MOI 0.01 for 2 hours, washed with PBS, and the medium replaced with DMEM containing 2% FBS. When cultures were fully infected (exhibited greater than 90% cytopathic effect (CPE)), flasks were freeze-thawed three times and supernatants collected and clarified at 3500 rpm for 30 minutes at 4 °C. Supernatant was collected, aliquoted and stored at −80°C. TCID_50_ was performed in VeroE6 cells in quadruplicate and infectious titre determined using the Reed-Muench method^20^. Clinical isolates representing SARS-CoV-2 variants including Alpha (CEPHR_IE_B.1.1.7_0221, GenBank accession ON350867, passage 2), Delta (CEPHR_IE_AY.50_0721, GenBank accession ON350967), Omicron (CEPHR_IE_BA.1_0212, GenBank accession ON350968, Passage 2) clinical isolates were isolated from SARS-CoV-2 positive nasopharyngeal swabs from the All-Ireland Infectious Disease (AIID) cohort (10). These were isolated and amplified on Vero E6/TMPRSS2 cells (#100978), obtained from the Centre For AIDS Reagents (CFAR) at the National Institute for Biological Standards and Control (NIBSC)(11).

### 2.4 SARS-CoV-2 infection and neutralisation

Cells were plated on collagen/poly-L-lysine coated 24-well plates, and 24 hours later, inoculated with SARS-CoV-2 for 1 hour at 37°C, washed, and incubated at 37°C. At the indicated times post-infection, cells were fixed for 1h with 4% paraformaldehyde (PFA) for immunofluorescent staining. Alternatively, to evaluate infectious viral titre within cells or released into the culture medium, supernatant was harvested and an equal quantity of culture medium added to cells which were freeze-thawed x3, centrifuged at 3,000rpm and supernatant collected from cellular lysate. Cellular supernatant (released virus) or lysate (intracellular virus) was titrated on VeroE6 cells. 72h post-titration, TCID50 was calculated according to the method of Reed and Muench, 1938 (12). To ensure that input virus was not present in supernatants, PBS from the final wash 1h post-infection was titrated on VeroE6 cells to confirm that no infectious virus was present.

To evaluate the ability of neutralising anti-ACE2 antibody to inhibit infection of human neural cells, 10 µg/mL antibody was incubated with the cells for 1 hour at 37°C prior to inoculation with SARS-CoV-2.

### 2.5 Immunofluorescent Staining and microscopy

Target cells (1 x 10^5^/mL) were plated on bovine type 1 collagen (BMEC), poly-L-lysine-coated Thermanox® coverslips (Nunc Thermo Fisher, Roskilde, Denmark), or 24 well cell culture plates, and incubated for 24 hours. Cells were exposed to SARS-CoV-2 as described in Section 2.4 and fixed with 4% PFA for 1h at 37°C. Primary antibodies anti-SARS-CoV-2 spike (Sino Biological, 20 µg/mL) and K1 anti-dsRNA (Scicons, 2 µg/mL) were incubated for 1h at room temperature, washed, and secondary antibodies Alexa 488/Alexa 594 (Invitrogen, 2 µg/mL) were incubated for 1h at room temperature. Cells were mounted with ProLong Gold containing 4’,6-diamidino-2-phenylindole (DAPI) counterstain, and imaged using an Olympus FV3000 confocal microscope or enumerated by manual counting using a Zeiss Axio Imager epifluorescent microscope. Confocal data were collected as z-stacks, deconvolved and maximum intensity projected. Infectivity is expressed as focus forming units per millilitre of virus (FFU/mL).

### 2.6 Real-Time Quantitative Reverse-Transcription Polymerase Chain Reaction

Real-time quantitative reverse transcriptase polymerase chain reaction (RT-qPCR) assays were performed on an ABI 7500 Fast real-time PCR machine (Applied Biosystems, Waltham, MA, USA) in a total volume of 20 µl, containing 5 µl of template and TaqMan Fast Virus 1-step mastermix (Applied Biosystems). Primer sequences and concentrations and thermal cycling conditions for SARS-CoV-2 nucleocapsid 1 gene were as previously described (13). Forward primer: GACCCCAAAATCAGCGAAT, reverse primer: TCTGGTACTGCCAGTTGAATCTG, probe:FAM-ACCCCGCATTACGTTTGGTGGACC-IBFQ. Primers and probes were all used at 500nM. Cycling conditions were RT (50 °C – 600 s), 95 °C – 30 s, 45 cycles (95 °C – 5 s, 60 °C – 30 s).

### 2.7 Statistical analysis

Statistical analyses were performed using Student’s *t* test in Prism 9.1.2 (GraphPad), with *P* < 0.05 being considered statistically significant and corrected for multiple comparisons when required (Bonferroni). Error bars show standard deviation and data are representative of three or more independent experiments with three replicates per experiment.

## 3. Results

### 3.1 Astrocytes, neurons and choroid plexus epithelium from the human central nervous system express ACE-2 and TMPRSS2

Frontal cortex and medulla from five healthy human donors were evaluated for expression of the SARS-CoV-2 entry receptor, ACE-2, together with TMPRSS2, which is reported to prime the spike protein of SARS-CoV-2 and is required for viral entry (Hoffmann et al., 2020). Expression of both receptors was restricted to astrocytes and neurons in both brain areas examined from all five donors (Figure 1). ACE-2 expression was predominantly observed in astrocytes, with expression also observed in neurons. In contrast, TMPRSS2 expression was predominant in neurons, with expression also observed in astrocytes. In a single donor for which choroid plexus was available, choroid plexus epithelium was strongly immunoreactive to both ACE-2 and TMPRSS2 (Figure 1). These data demonstrate that SARS-CoV-2 receptor ACE2, and TMPRSS2, is predominantly expressed in astrocytes, neurons and choroid plexus epithelium in normal human central nervous system.

**Figure 1:**
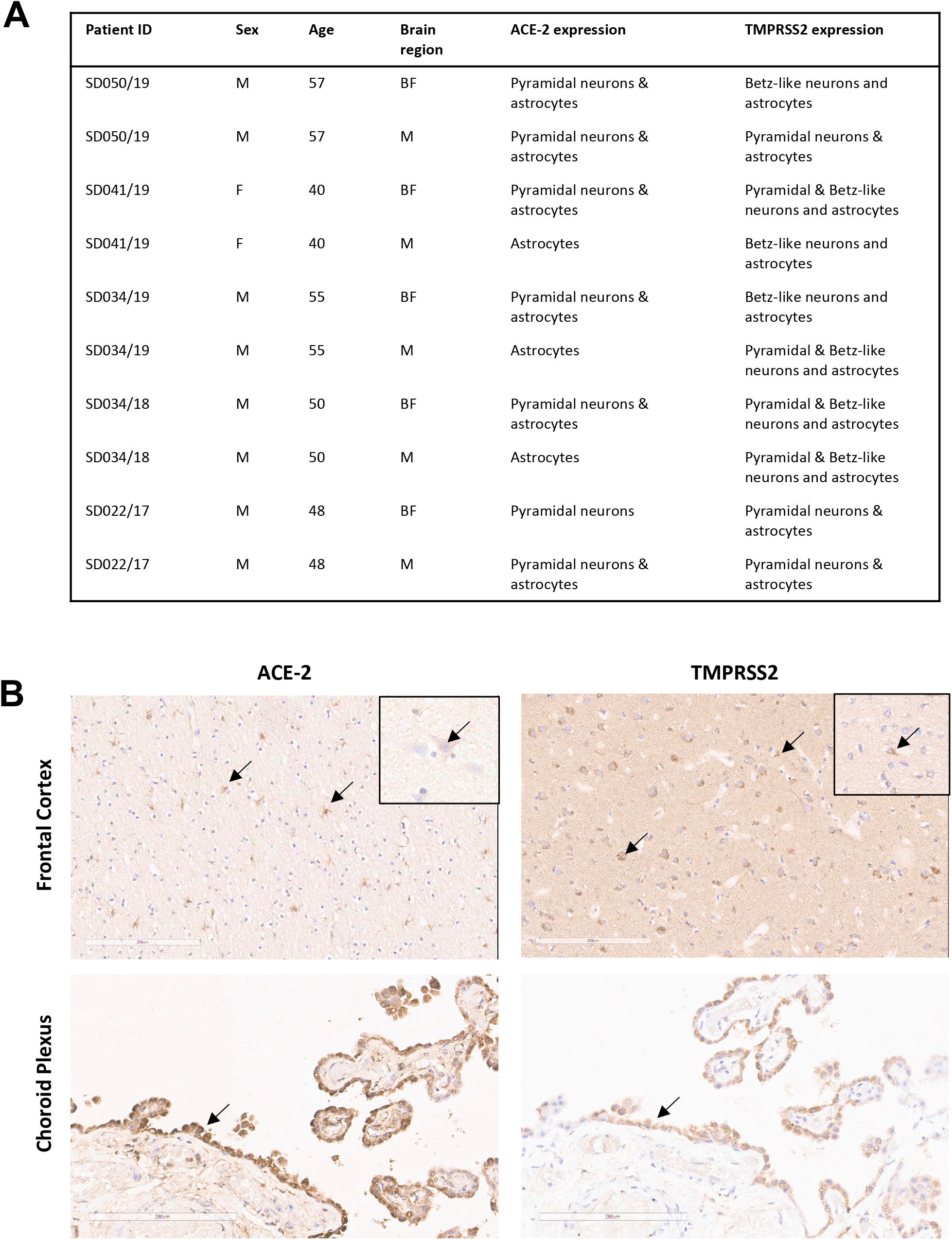
Human astrocytes, neurons and choroid plexus epithelium express ACE-2 and TMPRSS2. A) Sections of formalin fixed, paraffin embedded frontal cortex and medulla (10 µm) from five healthy donors were stained for immunoreactivity to ACE-2 and TMPRSS2. ACE-2 and TMPRSS2 expression was observed in astrocytes and pyramidal neurons within both brain regions, with predominantly astrocyte expression of ACE-2 and neuronal expression of TMPRSS2. Brain regions, BF: Medulla, M: Anterior frontal convexity. B) ACE-2 and TMPRSS2 expression in frontal cortex and choroid plexus reveals expression of ACE-2 in astrocytes, with expression also observed in neurons (insets). TMPRSS2 was predominantly observed in neurons with expression also observed in astrocytes (inset). Expression of both ACE-2 and TMPRSS2 was observed in choroid plexus epithelium (arrows). Bar −200µm.

### 3.2 SARS-CoV-2 infects human neurons, astrocytes, pericytes and choroid plexus epithelial cells

Primary human neurons, astrocytes, choroid plexus epithelial cells, brain microvascular endothelial cells (BMEC), brain vascular pericytes and immortalised human microglia were inoculated with SARS-CoV-2. Neurons, astrocytes and choroid plexus epithelial cells were found to support SARS-CoV-2 infection, with extremely low levels of infection observed in pericytes (Figure 2A). Astrocytes supported the highest levels of infection, with approximately 25-fold higher levels of infected cells than that of pericytes. Anti-SARS-CoV-2 spike and dsRNA staining was observed in infected cells (Figure 2B). In contrast, no infection of BMEC or microglia was observed. No cytopathic effect was observed in any cell type studied following SARS-CoV-2 inoculation, either by visual inspection or MTS assay (data not shown).

**Figure 2:**
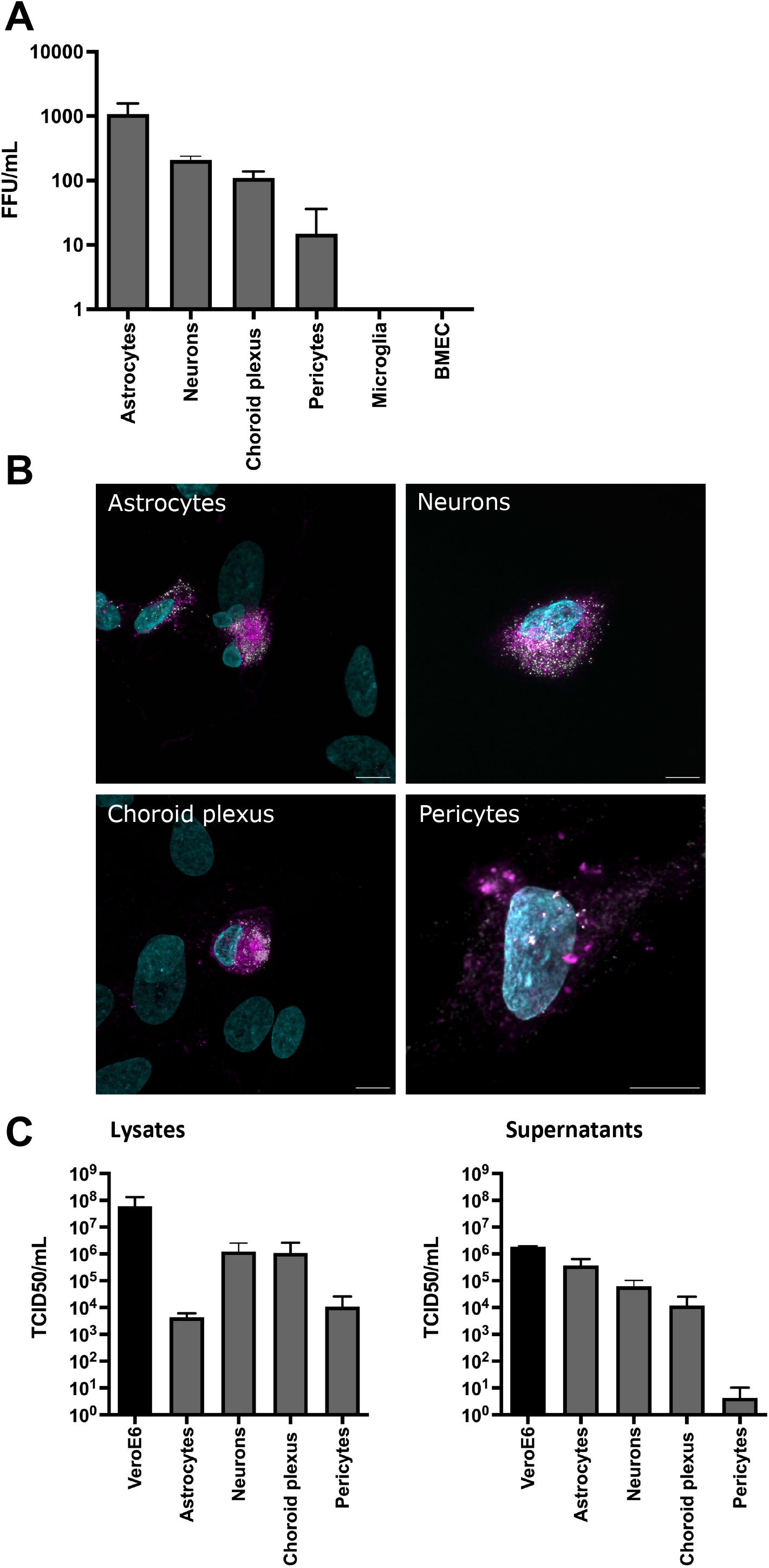
SARS-CoV-2 infects human neurons, astrocytes, pericytes and choroid plexus epithelial cells. (A) Primary human neurons, astrocytes, choroid plexus epithelial cells, brain vascular pericytes, microglia and brain microvascular endothelial cells (BMEC), were exposed to SARS-CoV-2 at an MOI of 0.1 for 1h. After 72h, cells were fixed and immunolabelled for anti-SARS-CoV-2 spike protein and dsRNA, and infected cells enumerated. Data are presented as focus-forming units/mL (FFU/mL). (B) Representative confocal images of SARS-CoV-2 infected cells. Anti-SARS-CoV-2 spike protein is shown in magenta, dsRNA (Scicons K1 mAb) in white, and DAPI in cyan. Scale bars: 10µm. (C) Cellular lysates and supernatants from primary human brain cells and VeroE6 cells were titrated on VeroE6 cells and 50% tissue culture infectious dose (TCID50) quantified after 72 h. N=3 independent experiments.

### 3.3 SARS-CoV-2 infected human neurons, astrocytes and choroid plexus epithelial cells support the full virus life cycle

To establish whether SARS-CoV-2 permissive brain cells were capable of supporting the full virus life cycle, infectious virus in the extracellular medium and cell lysate were quantified by titration on VeroE6 cells 72h post-infection of primary human brain cells. Primary human neurons, astrocytes, choroid plexus epithelial cells and pericytes supported the full virus life cycle, containing infectious virus within cells and released within cellular supernatants that was capable of infecting VeroE6 cells (Figure 2C). Neurons and choroid plexus epithelial cells displayed similar levels of intracellular infectious virus and virus released into the extracellular medium. In contrast, astrocytes released more infectious virus into the extracellular medium, with a titre of 3.675 x 10^5^/mL compared to 4.39 x 10^3^/mL for cell lysate (Figure 2C). Pericytes were permissive to infection, but released low levels of virus, with a 2500-fold decrease in virus titre in the supernatant compared to intracellular virus (Figure 2C). Infectious virus was not detected in the supernatants of cells following PBS washes post infection, confirming that virus detected in supernatants or cell lysates was not representative of input virus (data not shown). Furthermore, infectious virus was not detected from either cellular lysate or supernatant of BMEC or microglia.

### 3.4 SARS-CoV-2 infection of human brain cells is ACE2-dependent

To determine whether infection of primary human brain cells was ACE2-dependent, the effect of anti-ACE-2 neutralising antibodies on SARS-CoV-2 infection was investigated. In astrocytes and neurons, a significant decrease in infection in infection was observed compared to control infected cells. While a decrease in the number of infected choroid plexus epithelial cells was observed, this decrease was not statistically significant (Figure 3A). Due to the extremely low levels of SARS-CoV-2 infection in pericytes, it was not possible to evaluate anti-ACE-2 neutralisation in these cells. Using RT-qPCR to quantify SARS-CoV-2 N1 gene, a significant decrease in levels of N1 RNA in all three cell types following treatment with anti-ACE2 neutralising antibodies was observed (p < 0.0001)(Figure 3B).

**Figure 3:**
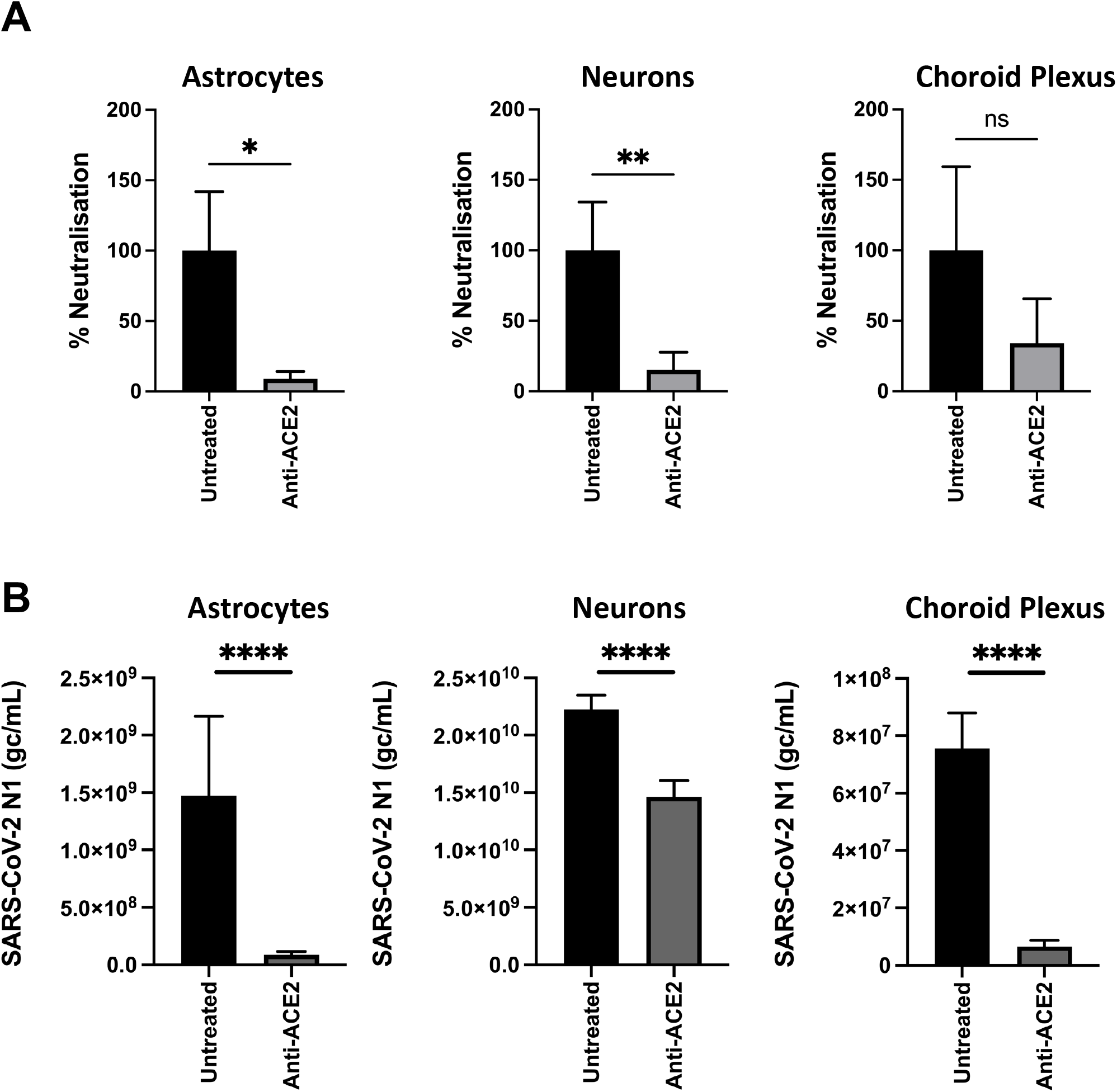
Primary human astrocytes, neurons and choroid plexus epithelial cells were incubated with ACE2 neutralizing antibody for 1h (10 μg/mL) before being inoculated with SARS-CoV-2 (Italy_INMI1) at an MOI of 0.1 for 1h. A) 72h post-infection, cells were fixed and immunolabelled for anti-SARS-CoV-2 spike protein, and infected cells enumerated. Data are presented as % neutralization relative to the untreated control. B) Cells were lysed and SARS-CoV-2 nucleocapsid (N1) gene quantified by qRT-PCR. Data are presented as SARS-CoV-2 N1 genome copies/mL (gc/mL). ****p<0.0001, **p<0.01, *p<0.05, ns=not significant. N=3 independent experiments.

### 3.5 SARS-CoV-2 variants infect human brain cells

Cells that were permissive to SARS-CoV-2 infection (neurons, astrocytes and choroid plexus epithelial cells) were infected with three clinical isolates of SARS-CoV-2 variants representing alpha, delta and omicron BA.1. Primary human neurons, astrocytes and choroid plexus epithelial cells supported infection by all three variants. Similar to our observations with Italy_INMI-1, BMEC and microglia did not support infection with SARS-CoV-2 variants. Pericytes were not infected with these variants, likely due to the lower titre of the clinical isolates compared with SARS-CoV-2 Italy_INMI-1, and therefore the lower likelihood of infection due to low permissivity of these cells to infection (data not shown). Neurons supported the highest levels of infection by all variants, and astrocytes and choroid plexus epithelial cells were infected at similar levels (Figure 4A). Representative images of SARS-CoV-2 infected cells revealed SARS-CoV-2 anti-spike immunoreactivity in infected cells (Figure 4B).

**Figure 4:**
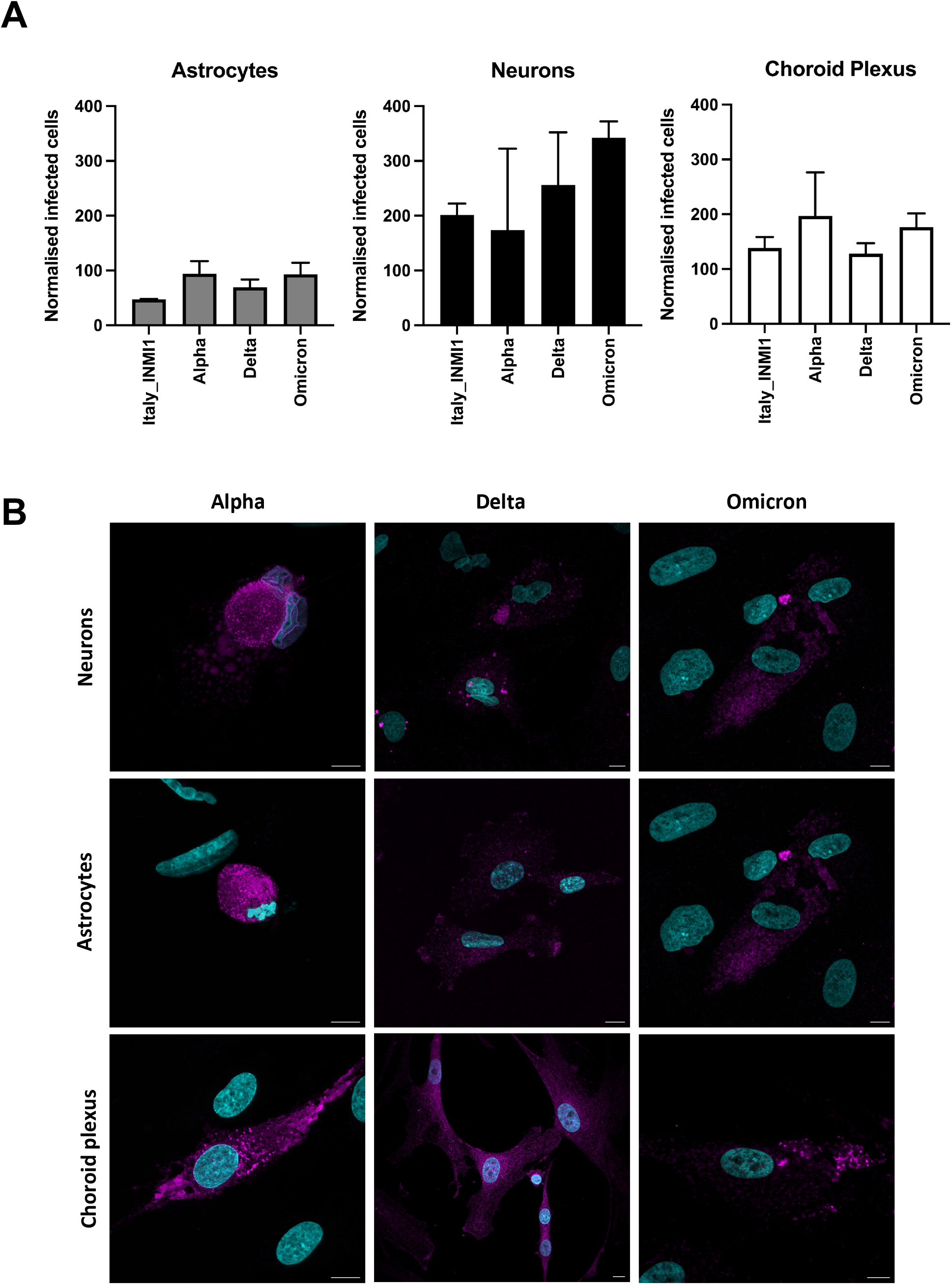
SARS-CoV-2 variants infect primary human brain cells. Primary human neurons, astrocytes, and choroid plexus epithelial cells were inoculated with SARS-CoV-2 variants (Italy_INMI1, Alpha, Delta and Omicron) at an MOI of 0.1 for 1h. After 72h, cells were fixed and immunolabelled for anti-SARS-CoV-2 spike protein and dsRNA, and infected cells enumerated. Data are presented as infected cell counts adjusted for virus titre and cell number. (B) Representative confocal images of SARS-CoV-2 infected cells. Anti-SARS-CoV-2 spike protein is shown in magenta and DAPI in cyan. Scale bars: 10µm.

## 4. Discussion

There is a growing body of evidence of neurological manifestations of COVID-19, and therefore, understanding the mechanisms of SARS-CoV-2 neuropathology will have a significant impact on treatment strategies. Neurological sequelae in COVID-19 patients are diverse, and include headache, encephalitis, vasculopathy and vasculitis including stroke, haemorrhage, and cerebral thrombosis (14). The underlying mechanism by which SARS-CoV-2, directly or indirectly, causes these manifestations is currently unclear and there may be multifactorial mechanisms. Moreover, peripheral nervous system sequelae including Gullain-Barre syndrome, peripheral neuropathy and myopathy, likely due to severe pulmonary disease, have been reported (14, 15). Recently, medium- and long-term follow up studies revealed an increased risk of dementia, migraine, Alzheimer’s disease and other cognitive defects 12-24 months post-infection. However, it is unclear whether SARS-CoV-2 exacerbated previously undiagnosed neurological disease, was associated with the Delta variant or confined to older age groups (16, 17).

SARS-CoV-2 binds to target cells at the initial stages of infection via the cellular receptor angiotensin converting enzyme-2 (ACE-2). SARS-CoV-2 binding to ACE-2 is followed by viral spike S’ subunit proteolytic cleavage at the cell surface by the transmembrane serine protease-2 (TMRPSS2)(18). In the present study, astrocytes, neurons and choroid plexus epithelial cells were the major cell types within normal human brain that expressed these SARS-CoV-2 entry factors. Similar profiles of receptor mRNA and antigen expression were observed in human brainstem in previous studies (19, 20). A recent, large scale imaging study of central nervous system pathological changes in COVID-19 patients revealed a loss of grey matter in several cortical areas including the limbic areas linked to the olfactory and gustatory system, providing a potential mechanism for the loss of taste and smell experienced by some individuals during acute infection (21). Moreover, a post-mortem study investigating SARS-CoV-2 antigen expression in the CNS of COVID-19 patients identified viral antigen in olfactory mucosa, in particular olfactory epithelial cells and neurons, which supports a model by which SARS-CoV-2 may invade the brain via the olfactory mucosa and the olfactory tract of the CNS through the cribriform plate (22).

The present study revealed that primary human primary neurons, astrocytes, pericytes and choroid plexus epithelial cells are permissive to SARS-CoV-2 infection and support the full virus life cycle *in vitro.* Several studies have used a range of neuronal cell lines, organoids and iPS cells to investigate the tropism of SARS-CoV-2 for the CNS, and have confirmed viral infection of these models, with disagreement as to which cell types support infection. Torices *et al.* (23) demonstrated that of cells of the neurovascular unit, astrocytes and microglia had the highest expression of SARS-CoV-2 receptors, and demonstrated that the human neuroblastoma SH-SY5Y cell line supported SARS-CoV-2 infection. In contrast, Pellegrini *et al.*, (5) demonstrated that SARS-CoV-2 does not infect neuronal cells within brain organoids, but instead cells of the choroid plexus epithelium; while Ramani *et al.,* (24) demonstrated preferential infection of neurons within brain organoids. Using iPSC-derived neurons and astrocytes, Kettunen *et al* observed that SARS-CoV-2 infected neurons at a low level, but not astrocytes (25). Few studies have investigated SARS-CoV-2 infection of primary, differentiated human cells. In agreement with the current study, Proust *et al* (26) demonstrated SARS-CoV-2 infection of primary human pericytes and astrocytes, with cell death in pericytes, which was not observed in the present study. While few studies have reported the presence of viral RNA in brain tissue from patients with COVID-19, a recent study demonstrated viral RNA and SARS-CoV-2 spike expression in 5 patients with histological changes suggestive of SARS-CoV-2 neuropathology. The main cell type expressing viral antigen was astrocytes and neurons, in agreement with the present study (26).

Various mechanisms of neuroinvasion by SARS-CoV-2 have been proposed, including viral disruption and invasion across the blood-brain barrier (BBB), blood-cerebrospinal fluid (CSF) barrier, via cranial nerves, including the olfactory nerve or trigeminal nerve, or anterograde or retrograde axonal transport (reviewed by (27)). The role of these potential mechanisms in SARS-CoV-2 neuroinvasion is currently unclear, and warrants further investigation. The central role of astrocytes in maintaining BBB integrity is of potential significance given our observation, in agreement with others, that astrocytes are potentially a significant target of SARS-CoV-2 replication. Disruption to astrocyte homeostasis could lead to BBB breakdown and viral and inflammatory mediator entry to the CNS, particularly if other mechanisms of viral entry play a role, such as viral invasion via cranial nerves. Similarly, choroid plexus epithelial cells form tight juntions and maintain blood-CSF barrier function, so disruption of this barrier opens a potential further portal of viral entry to the brain. Many therapeutics, such as those used as HIV antivirals, do not readily cross the BBB and reach therapeutic concentrations within the CNS (28). While further studies investigating the ability of SARS-CoV-2 to replicate in the brain *in vivo* are warranted, therapeutic strategies should consider the ability of antivirals to penetrate the brain in therapeutic concentrations.

In conclusion, we have identified astrocytes, neurons and choroid plexus epithelial cells as the main cell types within the human central nervous system to express the SARS-CoV-2 entry receptor ACE-2 and TMPRSS2. Furthermore, using primary human brain cells cultured *in vitro,* we confirmed that astrocytes, neurons, choroid plexus epithelial cells and pericytes support the full virus lifecycle and release infectious virions. In addition to ancestral SARS-CoV-2, clinical isolates of alpha, delta and omicron infected astrocytes, neurons and choroid plexus epithelial cells. This data supports a model whereby SARS-CoV-2 is neurotropic, and further studies investigating the *in vivo* relevance of these data, and studies investigating the implications for CNS homeostasis and cerebral barrier integrity in SARS-CoV-2 infected individuals are warranted.

## Acknowledgements

This work was funded by a Science Foundation Ireland COVID-19 Rapid Response grant (20/COV/8492). The funders had no role in study design, data collection and interpretation, or the decision to submit the work for publication. The authors acknowledge the University College Dublin Veterinary Medicine Biosafety Level 3 laboratory for facilitating experiments with SARS-CoV-2.

